# Expansion of the circadian transcriptome in *Brassica rapa* and genome-wide diversification of paralog expression patterns

**DOI:** 10.1101/2020.04.03.024281

**Authors:** Kathleen Greenham, Ryan C. Sartor, Stevan Zorich, Ping Lou, Todd C. Mockler, C. Robertson McClung

## Abstract

An important challenge of crop improvement strategies is assigning function to paralogs in polyploid crops. Gene expression is one method for determining the activity of paralogs; however, the majority of transcript abundance data represents a static point that does not consider the spatial and temporal dynamics of the transcriptome. Studies in Arabidopsis have estimated up to 90% of the transcriptome to be under diel or circadian control depending on the condition. As a result, time of day effects on the transcriptome have major implications on how we characterize gene activity. In this study, we aimed to resolve the circadian transcriptome in the polyploid crop *Brassica rapa* and explore the fate of multicopy orthologs of Arabidopsis circadian regulated genes. We performed a high-resolution time course study with 2 h sampling density to capture the genes under circadian control. Strikingly, more than two-thirds of expressed genes exhibited rhythmicity indicative of circadian regulation. To compare the expression patterns of paralogous genes, we developed a program in R called DiPALM (Differential Pattern Analysis by Linear Models) that analyzes time course data to identify transcripts with significant pattern differences. Using DiPALM, we identified genome-wide divergence of expression patterns among retained paralogs. Cross-comparison with a previously generated diel drought experiment in *B. rapa* revealed evidence for differential drought response for these diverging paralog pairs. Using gene regulatory network models we compared transcription factor targets between *B. rapa* and Arabidopsis circadian networks to reveal additional evidence for divergence in expression between *B. rapa* paralogs that may be driven in part by variation in conserved non coding sequences. These findings provide new insight into the rapid expansion and divergence of the transcriptional network in a polyploid crop and offer a new method for assessing paralog activity at the transcript level.

**Significance:** The circadian regulation of the transcriptome leads to time of day changes in gene expression that coordinates environmental conditions with physiological responses. *Brassica rapa*, a morphologically diverse crop species, has undergone whole genome triplication since diverging from Arabidopsis resulting in an expansion of gene copy number. To examine how this expansion has influenced the circadian transcriptome we developed a new method for comparing gene expression patterns. This method facilitated the discovery of genome-wide expansion of expression patterns for genes present in multiple copies and divergence in temporal abiotic stress response. We find support for conserved sequences outside the gene body contributing to these expression pattern differences and ultimately generating new connections in the gene regulatory network.

## Introduction

The transition from basic research in Arabidopsis to new model systems for monocot and dicot crops has focused attention on the implications of polyploidy on our current models of genetic processes developed in Arabidopsis. The expansion of gene content through whole genome duplication (WGD), tandem duplication or transposed duplicates has been predicted to account for the evolution of morphological complexity (1). Improving crop yield in rapidly changing climates depends on our ability to integrate these gene content expansions into functional classifications of physiological importance. This will rely on the growing collection of sequenced genomes, not just across crop species but of ecotypes within species, including complementary genomic datasets such as transcriptomes, methylomes, chromatin accessibility and metabolomic profiling. One difficulty in assigning new or overlapping functions among paralogs arises from heterogeneity in transcript abundance datasets generated under various environmental conditions, from various tissue types, and at distinct times of day. Many studies have explored the potential for functional divergence of duplicated genes by comparing expression levels normalized across a collection of expression studies (2–5) which limits the search to genes showing very dramatic differences in transcript abundance at a single time point.

The importance of daily rhythms was recognized with the 2017 Nobel prize in physiology or medicine to Jeffrey Hall, Michael Rosbash and Mike Young for their discoveries of the molecular mechanisms generating circadian rhythms in Drosophila (6, 7). The conservation of circadian oscillators across the animal and plant lineages supports a role for these rhythms in maintaining fitness and evolving new regulatory pathways to fulfill that role (8). Many lines of evidence support the importance of circadian rhythms to plant biology, including photosynthesis, starch metabolism, biomass accumulation, and reproduction (9, 10). The circadian clock responds to environmental conditions to set these circadian rhythms to local time (10). As a consequence, circadian rhythms and thus much of plant biology are likely to be influenced by climate change. Examples of natural variation in plant circadian function are accumulating, as is evidence that many domestication traits that facilitated the geographic expansion of crops are due to alterations in circadian clock genes (11–13). This supports the utility in targeting circadian clock processes as a means of trait improvement without disrupting critical pathways required for growth and yield.

Plant circadian biologists have focused primarily on Arabidopsis as a model for defining circadian clock components and function in plants (14). Transcriptome studies have revealed extensive circadian control of transcript abundance resulting in time of day changes in expression (15–17). These rhythmic changes in transcript abundance are not unexpected given the daily changes in light, temperature and precipitation that affect physiological processes such as photosynthesis. Dynamic changes in metabolism and physiology must be driven by dynamic changes in gene expression and ultimately protein regulation and activity. To identify candidate circadian regulators for trait improvement in crops, more detailed time course resolution of transcript abundance levels is needed to confirm whether the diel and circadian patterns observed in Arabidopsis are maintained in highly polyploid crops. The crop plant *Brassica rapa* offers an excellent model system for studies in crops. It is a member of the Brassicaceae and close relative of Arabidopsis making comparative studies feasible. The morphological diversity in *B. rapa* with turnip, Chinese cabbage, pak choi, leafy and oil-type varieties provides a wealth of phenotypic traits to study in one species allowing for broad applicability to other crops. Preliminary studies have shown diversity in circadian clock parameters among morphotypes that correlate with various physiology measures suggesting that circadian clock variation has contributed to *B. rapa* diversification (18). Examination of the orthologs of known circadian clock genes in Arabidopsis revealed preferential retention of these genes in *B. rapa* following the triplication and extensive fractionation of the genome after diverging from Arabidopsis around 24 million years ago (MYA) (19). The preferential retention of clock genes suggests that their involvement in protein complexes and regulation of critical pathways makes them sensitive to dosage effects. The gene dosage balance hypothesis proposes that duplication of the entire genome is favored over single or chromosome level duplications because it maintains the appropriate concentration of gene products (20). This is supported by studies in yeast where genes of protein complexes tend to be lost simultaneously with their interacting proteins (21). The increase in expression of one duplicate could lead to or permit loss of the other duplicate or neo-functionalization (21).

To assess the functional significance of the retention of circadian clock genes in *B. rapa* and look for possible examples of neo-functionalization we performed two high resolution circadian transcriptome experiments to characterize the circadian network. To compare the expression dynamics of paralogous genes, we developed a novel method for identifying and classifying changes in expression patterns. This method is available as an R package called DiPALM (Differential Pattern Analysis via Linear Models). DiPALM facilitated a comparison of paralog expression patterns, revealing genome-wide expansion of phase domains among paralogs providing novel insight into the rapid divergence of the transcriptional network in this crop. We applied DiPALM to our recent drought time course experiment in *B. rapa* and discovered differential responses to mild drought stress among paralogs suggesting evidence for neo- and sub-functionalization. Using previously generated circadian microarray data in Arabidopsis we compared gene regulatory networks (GRNs) to identify the more Arabidopsis-like versus the more divergent (less Arabidopsis-like) among pairs of *B. rapa* paralogous Transcription Factors (TFs) based on conservation of connected targets in the network. The identification of the more Arabidopsis-like TF ortholog was supported by the presence of conserved noncoding sequences (CNSs) surrounding TF target genes, reinforcing the importance of these CNSs for regulating gene expression.

## Results

### How pervasive is circadian regulation of the transcriptome in *B. rapa*?

The preferential retention of genes contributing to circadian clock function in multiple copies in *B. rapa* (19) suggests that relative dosage of clock proteins is important. Have these retained paralogs diversified in function and contributed to robustness and flexibility in the circadian clock? If the clock were essential for plant growth and coordinating responses with the environment, we would expect that the circadian regulation of the transcriptome would be similarly impacted. To examine the extent of circadian regulation of the *B. rapa* transcriptome and the expression patterns of circadian regulated paralogs, we designed two RNA-seq experiments. The first photocycle (LD) experiment involved entraining *B. rapa* yellow sarson (R500) plants under 12 h light/ 12 h dark and constant 20°C for 15 days after sowing (DAS) before transfer to constant light and 20°C (LDHH). The second thermocycle (HC) experiment involved entraining *B. rapa* R500 plants under 24 h light with 12 h 20°C and 12 h 10°C temperature cycles (LLHC) until 15 DAS before transfer to LLHH. Following 24 h in constant conditions leaf tissue from the youngest leaf was collected and flash frozen in liquid nitrogen every 2 h for 48 h. These two conditions were designed to capture the genes under circadian regulation driven by light and temperature zeitgebers (German for “time givers”, referring to entraining signals). Studies in Arabidopsis have demonstrated widespread circadian regulation of the transcriptome with distinct and overlapping genes involved in various entraining conditions (16, 22).

To identify the circadian transcriptome, we analyzed the LD and HC datasets and ran the circadian analysis program RAIN (23), a nonparametric method for the detection of rhythms from a variety of waveforms that are typical of transcript abundance datasets. The 2 h sampling regimen provided the resolution to capture more cycling genes than possible with typical 4 h sampling (24). Using a Benjamini-Hochberg corrected p-value of 0.01, we identified 16,973 high confidence circadian regulated genes from the two datasets. Of the 22,204 genes that were expressed in the RNA-seq datasets, 76% of them passed our cutoff for cycling in one or both conditions, indicating retention of circadian regulation of the transcriptome following WGD in *B. rapa*. To assign cycling genes to specific phase bins based on timing of peak expression we generated co-expression networks for each dataset using the weighted gene correlation network analysis (WGCNA) package in R (25). Applying a network approach to time series data has proven to be an effective method for grouping similarly phased genes based on their expression pattern (26). This resulted in 14 modules in the LD dataset and 10 modules in the HC dataset. To demonstrate the uniformity of the genes within each module, a heat map was generated with the log2 transformed expression data of each gene for all modules numbered based on their phase (Fig. 1). From the heatmaps, it is evident that each module reflects a similarly phased set of genes that collectively are phased throughout the day with the LD_01 module showing peak expression at ZT24 (subjective dawn) and LD_09 showing peak expression at ZT36 (subjective dusk). Most striking is the proportion of the transcriptome that exhibits rhythmic patterns of expression with 13,474 genes in LD clustered into just 14 modules and 14,211 genes in HC clustered into just 10 modules; these data establish that a substantial portion of the transcriptome is under circadian regulation in *B. rapa*.

**Figure 1.**
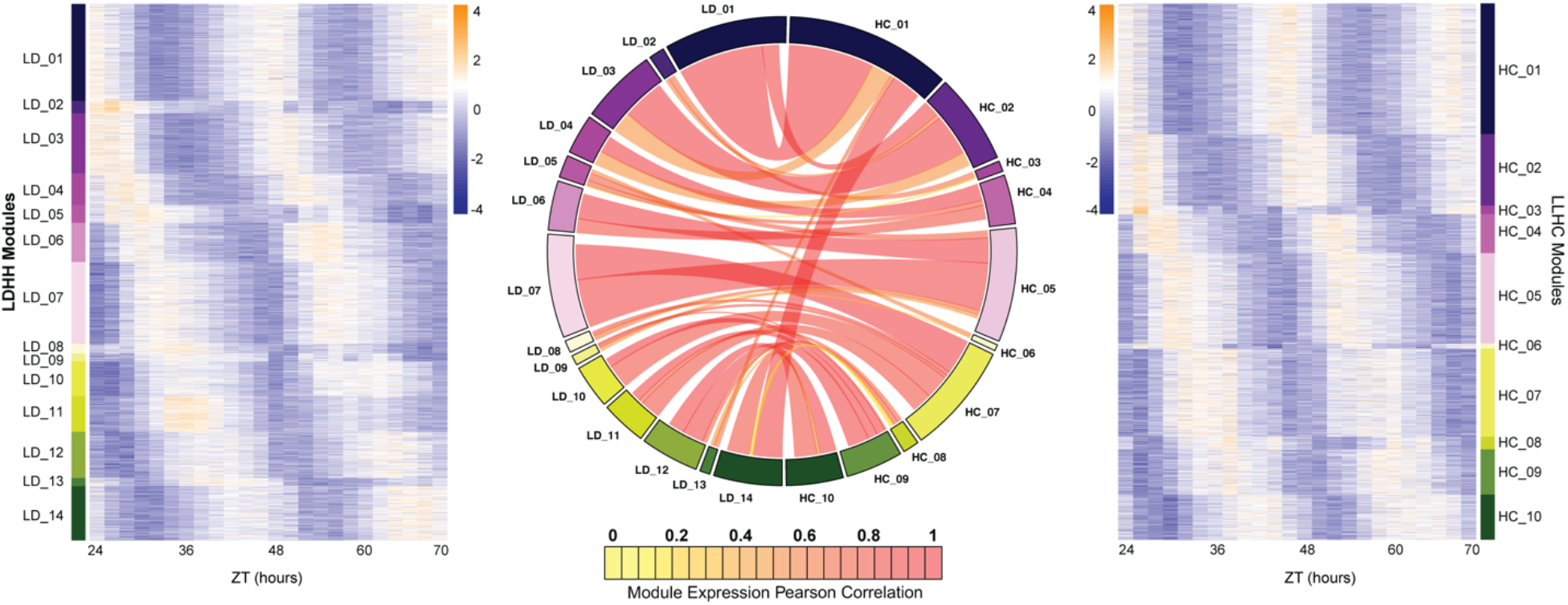
Expansion of the circadian transcriptome in *B. rapa*. Co-expression networks were generated for the LDHH (LD) and LLHC (HC) datasets. The 14 modules from the LD dataset are shown in the heatmap on the left and the 10 modules from the HC dataset are shown on the right. Heatmaps were generated using the log2 transformed FKPM expression data. Low expression level is in purple and high expression level in orange. The circos plot in the middle shows the overlap in genes between the LD and HC modules. Modules are numbered based on their phase starting at the beginning of the day (ZT24). The size of the ribbon signifies the number of genes in common between the connected modules and the color represents the Pearson correlation between the eigengenes of the two datasets.

The similarity in patterns seen in the LD and HC heatmaps in terms of phasing and distribution of genes within those phase bins suggests that there may be considerable overlap in gene phasing under LD and HC entrainment. To quantify the overlap, we matched the genes across the two networks and compared the correlation of eigengenes between LD and HC modules. The circos graph in Fig. 1 depicts the overlap between the two networks where the size of the ribbon represents the number of genes in common between the two modules and the color signifies the Pearson correlation between the eigengenes of the two datasets with dark orange being a correlation of 1. Because the modules are numbered based on their phase, similarly phased modules are arranged in the same order in the circos plot and the significant overlap and expression pattern between these modules is evident. This comparison demonstrates that most genes have the same or similar phasing when entrained by either photocycles or thermocycles. To further assess the similarity between these two datasets, GO ontologies were compared for each module from the LD and HC experiments (Dataset S1). This revealed similar biological processes enriched in the modules with significant correlation in expression pattern. For example, genes in LD_03 and HC_02 were both significantly enriched for photosynthetic processes and response to abiotic stress, consistent with their morning phased expression. Interestingly, genes in LD_07 and HC_07 were significantly enriched for protein phosphorylation suggesting a time-of-day dependency for the regulation of this process.

The correlation between module membership is not a rigorous test of differential transcript abundance and many genes have low correlations with their module eigengene that are not reflected in the analysis in Fig. 1. To our knowledge, there was not a rigorous test available for identifying significantly different gene expression patterns that would classify the change based on phase, amplitude or a combination of both. Rather than looking at differentially expressed genes at any given time point, we felt it was more important to classify a pattern change that encapsulated the entire time course. This led us to the development of the R package DiPALM. DiPALM takes advantage of the network analysis that assigns an eigengene to each module and therefore produces a minimal set of patterns representing the entire data set. The expression correlation of a given gene to any module’s eigengene defines the module membership (kME) of that gene to the module. The combination of kMEs for a gene across all modules can be used to encode its expression pattern numerically and allows for quantitative comparisons between any two gene’s expression patterns. This allowed us to run a set of linear model contrasts (one for each eigengene) that is analogous to running a contrast of gene expression data between time points or treatments except in this case the kME value represents the entire expression profile across the time course. We first tested for differential expression patterns between the LDHH and LLHC datasets. To generate a significance cutoff, we also ran the analysis on a permuted gene expression set where gene accessions were randomly re-assigned to expression patterns. P-values were then calculated using this permuted set. Using a p-value cutoff of 0.01, we identified just 1713 genes, or 11% of all cycling genes, that have altered patterns between LDHH and LLHC entrainment. To quantify overall expression level variation, we ran a similar linear model analysis on the median expression level for all genes and identified 3465 (23%) genes, only 448 of which overlapped with the pattern change list (Dataset S2). The 11% of cycling genes with entrainment-dependent cycling patterns are very interesting but further analysis in this area is not within the intended scope of this manuscript. A functional enrichment analysis of these genes revealed translation initiation factors and ncRNA metabolic process among the significant functional categories (Dataset S1, “Differential_Pattern” Tab). Given that the majority of genes show similar expression between the LD and HC datasets, we have combined the datasets for all further analysis in order to increase our statistical power by having four replicates per time point rather than two. One significant advantage of a linear model-based frame-work is the ability to account for any identified effect. Therefore, for all subsequent references to the RNA-seq dataset we combined LD and HC entrainments and included the LD/HC factor as a covariate in the linear model.

### Do retained multi-copy circadian regulated genes exhibit gene dosage behavior?

The network analysis revealed that a large portion of the transcriptome exhibits rhythmic expression patterns. This would imply that multi-copy paralogs have retained their rhythmic expression patterns and circadian regulation consistent with the preferential retention of circadian clock orthologs in *B. rapa* (19). These results provide the first evidence of genome-wide expansion and retention of circadian control of the transcriptome following a WGD in *B. rapa*. Based on the gene dosage model, the balance in expression among different subunits of protein complexes must be maintained resulting in the proper adjustment of paralog expression level, in some cases resulting in one copy maintaining high expression while the other is repressed (20). To evaluate this model, we calculated the mean expression level for all cycling genes across the 48 h time course. The mean expression levels for the set of retained multi-copy paralogs was significantly higher than for genes retained in single copy (Fig. 2*A*). It is possible that one of the retained copies is expressed at a much higher level than the average single copy gene as well as its paralog. To test whether this is the case, we separated all the cycling 2- and 3-copy paralogs into the highest and lowest expressed copies. We compare this to randomly paired single-copy gene pairs that were also separated into high and low groups (Fig. 2*B*). Surprisingly, the difference is observed in the low expressed paralog where these duplicated genes have significantly higher average expression levels compared to the genes retained in single copy. This suggests that although the duplicate paralogs do appear to exhibit gene dosage, their overall expression is retained at a higher level than expression of single copy genes.

**Figure 2.**
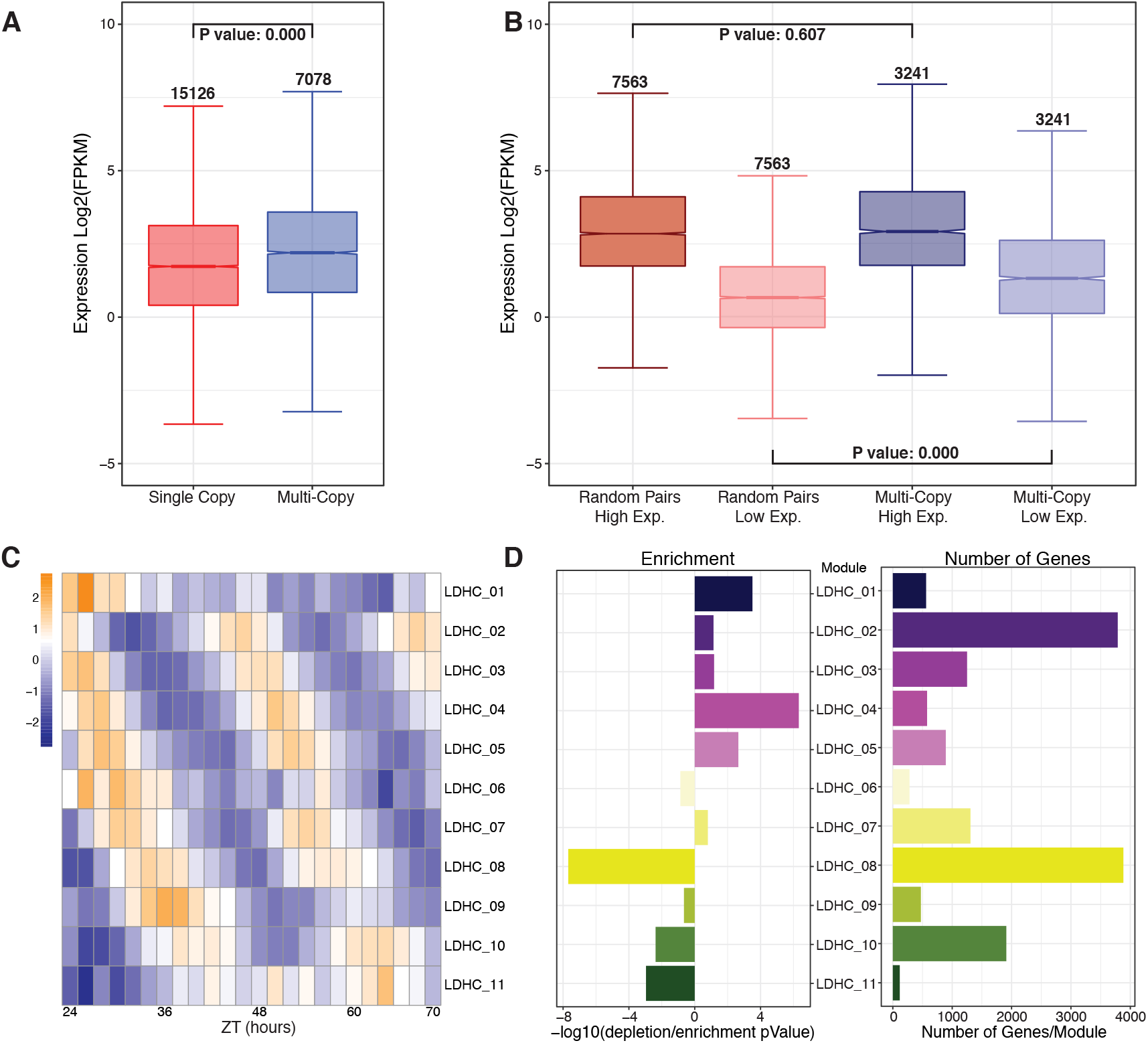
Retained multi-copy circadian regulated genes are highly expressed and display time of day variation. A. Mean log2 FPKM expression levels for each gene across the combined LD & HC time course. for multi-copy paralogs compared to single copy genes. Numbers above the whiskers indicate the number of genes in each group. P value is the result of an ANOVA test. B. Expression level comparison when paralogs are separated into high and low expression groups compared to randomly paired single copy genes. Numbers above the whiskers indicate the number of genes in the groups. P value is the result of an ANOVA test. C. Heatmap of the 11 modules of the combined LDHC co-expression network arranged by ZT time (in Zeitgeber [ZT] time, where ZT0 represents the most recent dark to light or cold to warm transition) across the x-axis and circadian phase along the y-axis. D. Results of a hypergeometric test of the number of multi-copy genes in each module. The left barplot shows the results of the hypergeometric test expressed as a −log10 P value with enrichment to the right and depletion to the left of 0. The right barplot shows the number of multi-copy genes in each of the modules.

The retention of multi-copy genes that are under circadian regulation and maintained at a relatively high expression level led us to explore whether there is evidence for divergence in expression pattern that would support neo- or subfunctionalization among paralogs. To associate similar patterns we applied the same WGCNA method to the combined dataset as was done for the individual analysis shown in Fig. 1. This resulted in 12 modules with distinct phasing throughout the day that is clearly visible when the eigengene expression for each module is presented as a heat map (Fig. 2*C*). We next wondered whether there was any association between phase of expression and retained copies that may suggest certain biological processes that are phased to specific times of day and may preferentially retain multi-copy genes. Based on the number of genes within each module, we ran a hypergeometric test to look for over- and under-enrichment of multi-copy genes within the modules. Surprisingly, we found that modules with phasing from morning to midday tend to be enriched for multi-copy genes. In contrast, evening and night phased modules were depleted for multi-copy genes (Fig. 2*D*). These trends were not associated with the number of genes within the module as can be seen with two of the largest modules LDHC_02 and LDHC_09 being over- and under-enriched, respectively. GO enrichment was carried out on a combined group of all multi-copy genes from the 5 morning modules with significant enrichment for multi-copy genes (p-value <0.05). The same was done for the group of all multi-copy genes from all 4 evening modules with significant depletion in multi-copy genes. Both of these sets appear to be representative of the whole modules from which they came with the morning-phase copied genes being significantly enriched for photosynthesis, translation and response to abiotic stimulus genes. The evening-phase multi-copy genes were significantly enriched for protein phosphorylation and glycosinolate biosynthesis genes, consistent with a phase at dusk (Fig. 2*D*, Dataset S3). Given the importance of proper transcriptional regulation of photosynthetic processes and the balance of enzyme components it is not surprising that there is a higher retention of multi-copy genes within these pathways. However, whether these genes are performing similar functions to their orthologous counterparts in Arabidopsis or have acquired new functions that could lead to additional regulation of the photosynthetic process is one exciting avenue of future study.

To look for signs of possible neo- or sub-functionalization among paralogs, we compared the expression profiles to identify paralogs with significantly different expression patterns. For 3-copy paralogs, where all three copies were expressed, these sets were converted into three, 2-copy pairs. We applied a similar linear modeling test using DiPALM that we ran on the LDHH and LLHC comparison but including a covariate in the model to account for differences in LD vs. HC. We ran this analysis on 4433 pairs where both genes are expressed. We found 3743 (84%) pairs with differential median expression (exDif) and 1883 (42%) paralog pairs exhibiting differential expression patterns (pDif) the vast majority (1607; 85%) of which overlapped with the exDif set (Dataset S4). Thus, 42% of expressed paralog pairs have diverged in circadian expression pattern in the R500 genome. However, this does not describe how the patterns differ. As with standard differential expression tests, it is critical to associate a direction of change in expression to know how a gene transcript is affected by a treatment or condition. To isolate the type of pattern change for the paralogous pairs exhibiting significantly different patterns, we performed clustering on the vector of expression values across the combined LD and HC data for each gene whose expression differed significantly from its paralog. This clustering had the effect of grouping genes based on their phase. Similar clustering was done for the significant exDif set. This clustering component is also part of the DiPALM package. A detailed description and example dataset of the analysis pipeline is provided with the package on CRAN (27).

To visualize the degree of pattern change, we generated a heat map of each of the clustering methods with the paralogous gene pairs stacked for comparison. As expected, the pDif clustering uncovered the changes in rhythmic patterns between pairs (Fig. 3*A*) while the exDif uncovered overall changes in transcript abundance (*SI Appendix*, Fig. S1). From the heatmap visualization it is clear that the majority of the pattern change among paralogous pairs is the result of a phase shift in peak expression. In some cases (Fig. 3*A* and *B*, clusters 4 and 12) the pairs are completely antiphase. This suggests genome-wide expansion of phase domains among retained paralogs. The expansion of expression domains is reminiscent of the *PSEUDO-RESPONSE REGULATOR* (*PRR)* and *REVEILLE (RVE)* families of circadian clock genes that were retained following WGD as well as tandem duplication events (28). The *PRR* genes in Arabidopsis have a temporally sequential expression pattern with *PRR9* expressed just after dawn followed by *PRR7, PRR5, PRR3* and finally *PRR1/TOC1* expressed in the evening (29). The PRR proteins appear to retain some common functions but their diverged expression patterns results in differential contributions to the circadian network (30–33). This also emphasizes the benefit of applying a network framework to the data that associates similarly regulated genes. A coordinated regulation must underlie these changes since we see paralogs with similar expression patterns having their paralogous pair diverging in expression together.

**Figure 3.**
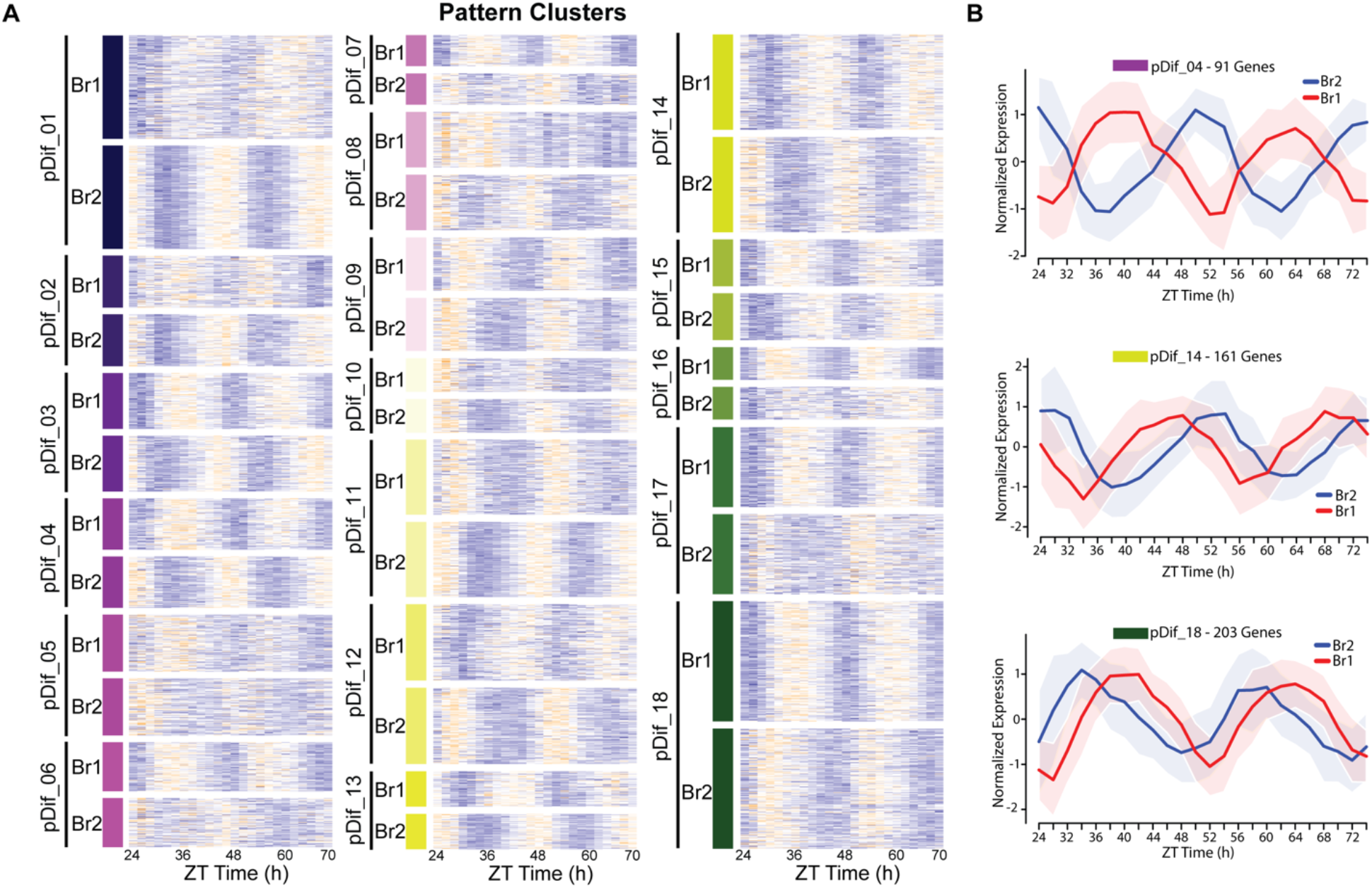
Divergence in phase domains among retained paralogs. A. Heat map of the results from the DiPALM pDif (A) clustering showing the changes in expression pattern for paralogous pairs. Each line of the heat map for each block corresponds to a paralogous pair. Three-copy paralogs were split into three 2-way comparisons. Expression values are log2 transformed FPKM values and the expression is arranged by ZT time across the x-axis. Higher expression levels are orange and low expression levels are purple. For example, pDif_01 shows paralogous pairs that are anti-phase with Paralog 1 peaking at ZT36 and ZT60 and Paralog 2 peaking at ZT24 and ZT48. B. Line plots showing the expression patterns for three modules (pDif_04, pDif_14, and pDif_18). Each plot shows the normalized expression of paralogous pairs for all the genes in the module. Ribbons (shaded regions) represent the standard deviation.

### Identifying the ‘Arabidopsis-like’ paralog of *B. rapa* using gene regulatory networks

The divergence in expression pattern among retained paralogs led us to speculate as to how diverged the retained pairs are with respect to their Arabidopsis ortholog (Fig. 4*A*). Are paralogous gene pairs equally likely to diverge in expression or does one copy retain the Arabidopsis-like expression pattern while the other copy acquires new expression variation? One method of comparing the orthologs between *B. rapa* and Arabidopsis is to compare phase of expression. However, assigning an accurate phase to circadian data from two cycles is challenging and often gene expression patterns show very broad peaks in abundance that can be difficult to classify, especially with the resolution of only 4 h that is available for Arabidopsis. Also, because we are comparing two species, we have to consider the properties of the clock in each species. The *B. rapa* R500 clock has a slightly shorter period than Arabidopsis Col-0 that results in apparently altered phasing among genes that is not indicative of divergence in function but arises merely from the different paces of the Arabidopsis and *B. rapa* oscillators.

**Figure 4.**
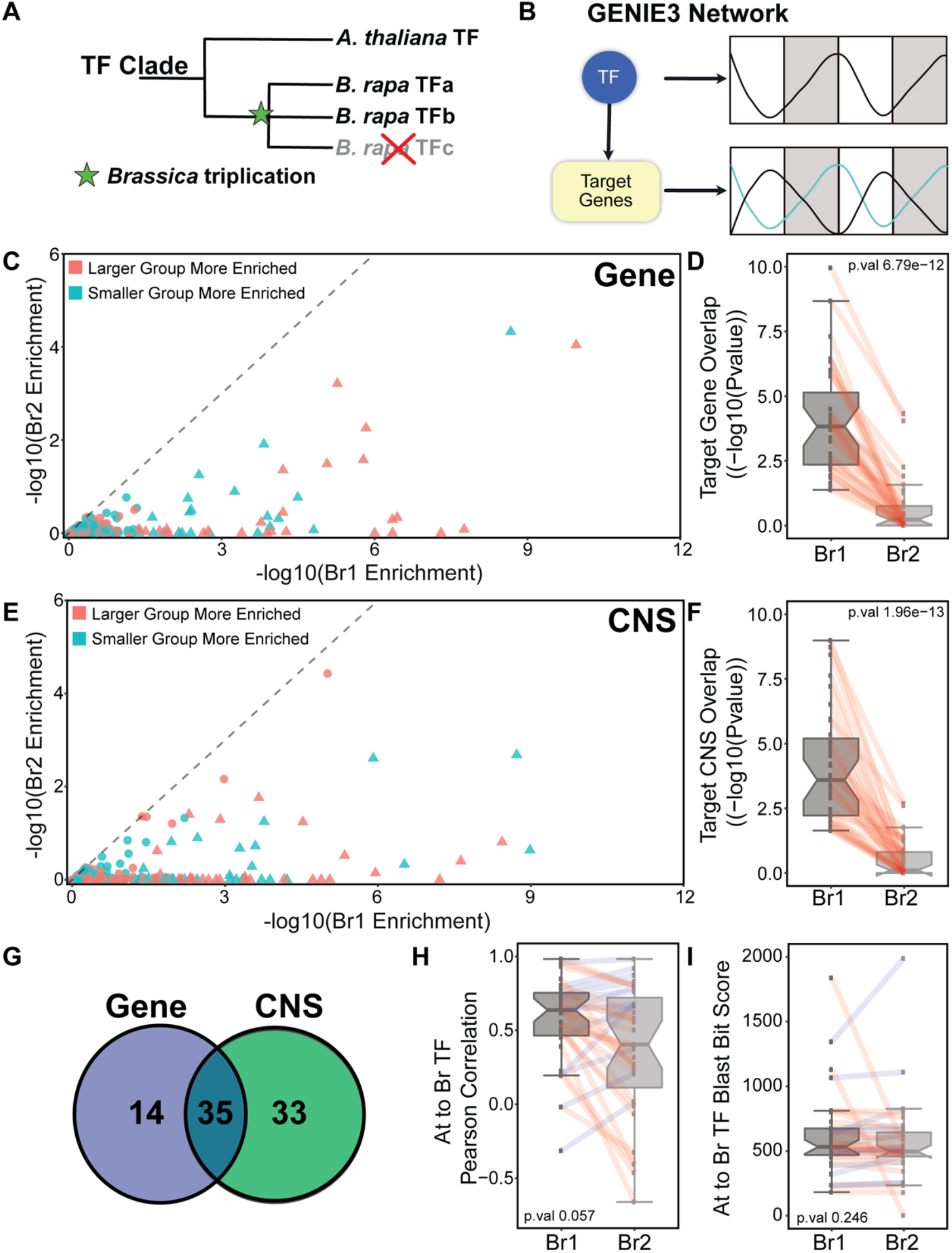
Identifying the ‘Arabidopsis-like’ paralog using GRNs. (A). Schematic showing the triplication event following the divergence between Arabidopsis and *B. rapa* leading to multi-copy orthologs of known Arabidopsis transcription factors (TF). (B) GENIE3 networks are generated to associate a TF with target genes based on the expression patterns of the TF and all the genes in the network. There is no assigned direction to the TF regulation in our GENIE3 network resulting in possible positive or negative regulation. (C&E) Scatterplots show all 256 *B. rapa* TF pairs in the analysis. Triangles indicate that they were identified as having one *B. rapa* TF significantly more Arabidopsis-like than the other; circles were not significantly different. Points colored red means that Br1 (the more Arabidopsis-like TF) has a larger target group and teal means Br1 has the smaller target group. (C) Shows the results for the gene overlap network analysis and (E) shows the results from the CNS overlap analysis. P value distributions of the overlap in *B. rapa* vs. Arabidopsis target genes (D) and CNSs (F). All Br1 and Br2 TFs are viewed as separate groups with paralogs connected by red lines. (G) Of the 49 TFs identified as more Arabidopsis-like based on target genes between the Arabidopsis and *B. rapa* networks, 35 overlapped with the 68 TFs identified based on CNSs between Arabidopsis and *B. rapa* target groups. (H) Distribution of the Pearson correlation of TF expression patterns comparing each Br1 and Br2 TF with Arabidopsis TF ortholog. (I) Distribution of BLAST bit score of Br1 and Br2 TFs compared to their orthologous Arabidopsis TF.

For example, if we plot the expression of single copy circadian clock genes from *B. rapa* and their corresponding orthologs in Arabidopsis we see a leading phase in *B. rapa* (*SI Appendix*, Fig. S2). To avoid these phase complications, we chose to use a gene regulatory network (GRN) approach that would provide additional statistical robustness by first predicting transcription factor (TF) targets based on expression dynamics within each species followed by a comparison of network connections between the species. We constructed the GRNs using GENIE3 (34). GENIE3 uses a random forest based method for GRN construction that demonstrated superior performance in the DREAM4 *In Silico Multifactorial* (35) and DREAM5 network inference (36) challenges (Fig. 4*B*).

We obtained previously published circadian microarray data from Arabidopsis that were generated under similar conditions with LD and HC entrainment (LL_LDHC, LL_LLHC, LL12_LDHH and LL23_LDHH from (17)). We selected the Arabidopsis TFs from the Arabidopsis TF database (https://agris-knowledgebase.org/AtTFDB/) and *B. rapa* TFs selected from the Mapman annotation “RNA regulation of transcription” in addition to known circadian clock TFs not included in the list. This resulted in a list of 612 Arabidopsis and 2147 *B. rapa* TFs that were expressed in their respective datasets. For the target set, we included 9201 Arabidopsis expressed genes and the corresponding 14,541 *B. rapa* expressed orthologs. Separate GRNs were generated for Arabidopsis and *B. rapa*. To identify significance of TF-target edges, we generated a permuted network by using the same TFs but shuffling the expression values for the target genes resulting in an Arabidopsis GRN with 71,216 edges and a *B. rapa* GRN with 947,062 edges. A gene was said to be a target of a TF if the edge between them was significant. We hypothesized that the paralogous TF in *B. rapa* that retained more of the Arabidopsis orthologous function would have a greater overlap in targets in the network compared to the more diverging pair. In other words, using TFs as features to describe the targets, how well can the expression of the target genes be explained by the expression of that TF (Fig. 4*B*)? A set of 256 TFs exist where one Arabidopsis TF can be associated with two *B. rapa* paralogs and all three of these genes had target groups defined by their respective GRNs.

For each of these 256 sets, we examined the significance of the overlap between the Arabidopsis TF target group vs. the corresponding *B. rapa* orthologous TF target groups. This resulted in two p-values for each group indicating how similar each *B. rapa* TF is to its orthologous Arabidopsis TF in terms of target gene overlap. Next we wanted to determine if the difference in these two p-values was significant; that is, does one of the *B. rapa* TFs show more conservation of target gene overlap with its Arabidopsis ortholog than the other *B. rapa* TF? This was accomplished with another permutation-based test where genes were randomly sampled to create target groups of the same sizes. The two *B. rapa* vs. Arabidopsis p-values were calculated and the difference was taken. This was repeated 10,000 times for each of the 256 TF sets. As a result, 49 TF pairs exhibited significant enrichment for one *B. rapa* TF (assigned Br1) being more similar to Arabidopsis than its paralog suggesting possible divergence in function between these TFs (Fig. 4*C* and *D*). It is worth noting that the size of the target group is not driving the enrichment as we see a broad distribution in target size and significance (Fig. 4*C*). Among the list of 49 TFs, six are part of the core circadian clock (*ELF3, ELF4, PRR9, PRR7, PRR5*, and *TOC1*), and a seventh, *RVE1*, integrates the circadian clock and auxin pathways (37, 38). Based on the predicted targets of these TFs in the GENIE3 model, there are 11,559 *B. rapa* and 3387 Arabidopsis genes regulated by these 49 TFs, providing further support for the retention and novel innovation of the circadian network in *B. rapa*. The divergence in TF target genes indicates several possible changes have occurred; these could include modifications to regulatory elements of the target genes, mutations that alter the TF protein binding efficiencies for motifs or interacting partners or a combination of both. Alterations to regulatory elements associated with core TFs and/or target genes can lead to whole pathway level restructuring. One possible mechanism for altered expression regulation is the distribution of conserved noncoding sequences (CNSs). A set of CNSs were identified across the Brassicaceae that show signs of selection (39). To associate CNSs with the R500 genome, we performed a BLAST analysis with a collection of ~63,000 CNSs against the *B. rapa* R500 genome. Provided the alignment met our BLAST filters, we allowed each CNS to have a maximum of three targets (see *Materials and Methods*). We repeated the BLAST with the Arabidopsis genome but restricted each CNS to one target gene.

To test for altered regulatory element occurrences between target genes of the identified diverging TFs we asked whether variation in CNS retention followed the same pattern as the observed gene expression changes. Do we see a similar divergence in *B. rapa* paralogous TF enrichment with Arabidopsis in the GENIE3 network if we replace the target set gene expression data with CNSs? With the list of genes and associated CNSs we replaced the target genes in the GENIE3 networks with CNSs resulting in a network with TFs targeting a group of CNSs rather than genes. We performed the same permutation tests to assign significant enrichment to the groups to ask whether we could identify a R500 TF ortholog that was more Arabidopsis-like than its paralogous pair. We identified 68 significant TFs (Fig. 4*E* and *F*), 35 of which overlapped with the 49 TFs identified based on target gene overlap (Fig. 4*G*). The agreement between these two approaches is apparent when the corresponding p-value distributions from the overlap of targets are plotted (Fig. 4*D* and *F*). In these boxplots, the Br1 enrichment for At target gene overlap is shown with the paralogous pairs connected by the red lines. Not only does the agreement between the target gene and CNS overlap further strengthen the support for those 35 TF pairs showing signs of divergence but suggests that the CNS distribution is associated with gene expression patterns and is a good predictor of expression variation. With this set of high confidence diverging TFs from the overlap group, we wondered whether changes in amino acid sequence contribute to the divergence between paralogous pairs in which case we would expect the Arabidopsis-like *B. rapa* TF to be more similar than its pair. To test this, we ran a protein BLAST using the *B. rapa* TFs against the Arabidopsis genome and examined the distribution of blast scores for the more and less Arabidopsis-like TF. Results from the BLAST suggest very little association between amino acid sequence and TF divergence (Fig. 4*I*) suggesting that changes to regulatory regions associated with target genes is likely to be a major driver of TF divergence. TF expression pattern changes are likely contributing to the divergence in regulation. To test this further, we conducted a similar analysis to the BLAST comparison where we looked at the expression correlation of *B. rapa* TFs vs. the orthologous Arabidopsis TF. In general, the more conserved *B. rapa* TF was more likely to maintain higher expression correlation to the Arabidopsis ortholog but the effect is not significant (Fig. 4*H*, p-value 0.057) and several of the 35 TF sets tested do not show this result.

However, due to the Arabidopsis and *B. rapa* datasets being slightly out of phase it is difficult to compare expression patterns directly (*SI Appendix*, Fig. S2). While paralogous TFs likely retain binding to the same motifs, a divergence in expression may result in the loss of transcriptional coactivators or corepressors required for gene activation or repression due to temporal separation in expression pattern resulting from the shift in phase of that TF. Similarly, an altered phase of expression might allow interaction with new TF interacting factors to provide new target affinity for that TF resulting in a new target set.

If the CNSs are driving the expression differences we would expect them to be enriched for TF binding motifs (39). To test for motif enrichment, we selected the overlap between the top 15 most significant TFs from the gene expression and CNS GRNs. This resulted in a list of 12 TFs that included those encoded by the circadian clock genes *TOC1* and *PRR5*. For each TF (*B. rapa* paralogs and Arabidopsis ortholog), we took the collection of CNSs represented by their target genes in the GRNs and ran them against the HOMER motif analysis algorithm (40). Because the CNSs are located throughout the gene (promoter, 5’UTR, introns, 3’UTR) we selected the sequence from 2kb upstream of the start codon to the 3’UTR for each target gene for comparison. We also included just the 2kb upstream sequence to compare to standard motif search parameters. For all 12 TFs tested, we found 3-5 fold greater enrichment for motifs in the CNS elements compared to the full length and 2kb promoter background sets (*SI Appendix*, Fig. S3). This is consistent with the results from the GRN analysis showing that CNSs are as predictive as expression dynamics and contain important regulatory elements. Further studies are needed to look for associations between groups of CNSs with their corresponding binding motifs and specific gene expression patterns. Since the GENIE3 algorithm associates target genes based on a TF being an activator or repressor, the target genes typically have several major expression patterns (Fig. 4*B*). Isolating distinct patterns and analyzing CNS variation between the target groups may reveal new motif groupings or novel motifs.

### *B. rapa* paralog expression pattern response to abiotic stress

In agreement with the gene balance hypothesis, which posits that multi-subunit complexes are sensitive to variations in stoichiometry resulting in dosage compensation to produce the same amount of product (41), the exDif clustering did reveal a consistent trend with one paralog having significantly higher median expression levels compared to the other retained paralog of that pair (*SI Appendix*, Fig. S1). However, as previously demonstrated, the overall expression levels for multi-copy genes is higher than the single-copy genes (Fig. 2*A* and *B*). In addition, the rhythmicity in the paralogous pairs is still apparent providing further support for focusing on the importance of the pattern of expression rather than simply the overall levels. This led us to wonder whether there is any indication that these pattern changes may contribute to new temporal responses to environmental stimuli such as abiotic stress. The gated stress response has been characterized in several plant species including Arabidopsis, poplar, rice and *B. rapa (26*, *42*–*45*). If a time-of-day dependent stress responsive gene in Arabidopsis now has two copies in *B. rapa* with altered expression patterns does this result in an expanded stress response window or does one copy retain stress response while the other loses it?

To look for indications of divergence in function we used our previous mild drought time course RNA-seq dataset (26) to test for altered responses to drought among pairs of paralogs. We first used the well-watered control samples to identify the paralogous pairs with altered patterns. Consistent with the divergence in pattern change under circadian conditions, the same trend is apparent under diel conditions. Out of 4664 total pairs where two copies show detectable expression levels, 3259 pairs had significantly different patterns under control conditions. In the circadian dataset we observed just 42% of genes with altered pattern but these diel data reveals 70% of pairs with altered pattern. Similarly, 77% of pairs (3602) had significantly different median expression levels (Dataset S5). Of the total pairs, 35% and 50% had one or both genes identified as circadian-regulated, respectively, indicating that both circadian and diel regulation drives the rhythms observed under diel conditions. For the pairs with altered patterns under the well-watered conditions, 93% were tested (both genes were expressed) under circadian conditions and 52% showed significant pattern changes.

To identify differential response to drought among paralogous pairs, we first ran DiPALM on all (23,248) expressed genes and identified 3891 with a significant pattern change and only 327 with median expression changes (Dataset S6). Unlike the circadian dataset, the largest source of variation in expression is due to a phase change consistent with the importance of time-of-day gating of stress response (26, 43). This provides further support for the dynamic nature of expression regulation and the limitations of simply quantifying transcript abundance differences at single time points. It should also be noted that this drought treatment captured the early signs of drought perception with very subtle expression changes during the first 24 h and more evident changes in the subsequent 24 h (26). The ability to detect these distinct patterns using DiPALM highlights the effectiveness of the network pattern approach for capturing unique and unpredictable patterns. To test whether the paralogous pairs exhibiting differential expression patterns under well-watered conditions are enriched for drought responsive genes we performed a permutation test. We randomly sampled the same number of pairs from the full set of pairs (3259 out of 4664) for 10,000 permutations to identify the likelihood of selecting drought responsive genes within the 3891 sampled set. As a result, the differentially patterned copies were enriched for genes with drought responsive patterns (P-value 0.0005) (Fig. 5*A*) whereas copies with differential median expression level were not enriched (P-value 0.6457) for drought responsive patterns (Fig. 5*A*). With enrichment of drought responsive genes among these pairs exhibiting different patterns, we wondered whether these pairs are more or less likely to have one or both copies responding to drought. Using the same 10,000 randomly sampled sets of 3891 pairs, we estimated a null distribution of the expected number of pairs with one and two drought-responsive genes. Results from the permutations indicated significant enrichment for pairs in which one member is drought responsive (P-value < 0.0001) and no enrichment for both copies being drought responsive (P-value 0.1852) (Fig. 5*B* and *C*). Thus, we observed enrichment for only one but not both paralogs responding to drought stress in *B. rapa*. This suggests that the genome-wide expansion of expression domains among paralogs is biologically meaningful, in this case for drought stress response. More broadly, these results have important implications for how we capture and characterize transcriptomic responses or ‘states’ when making predictions about paralog function. Temporal, spatial and conditional regulation can reveal new expression dynamics.

**Figure 5.**
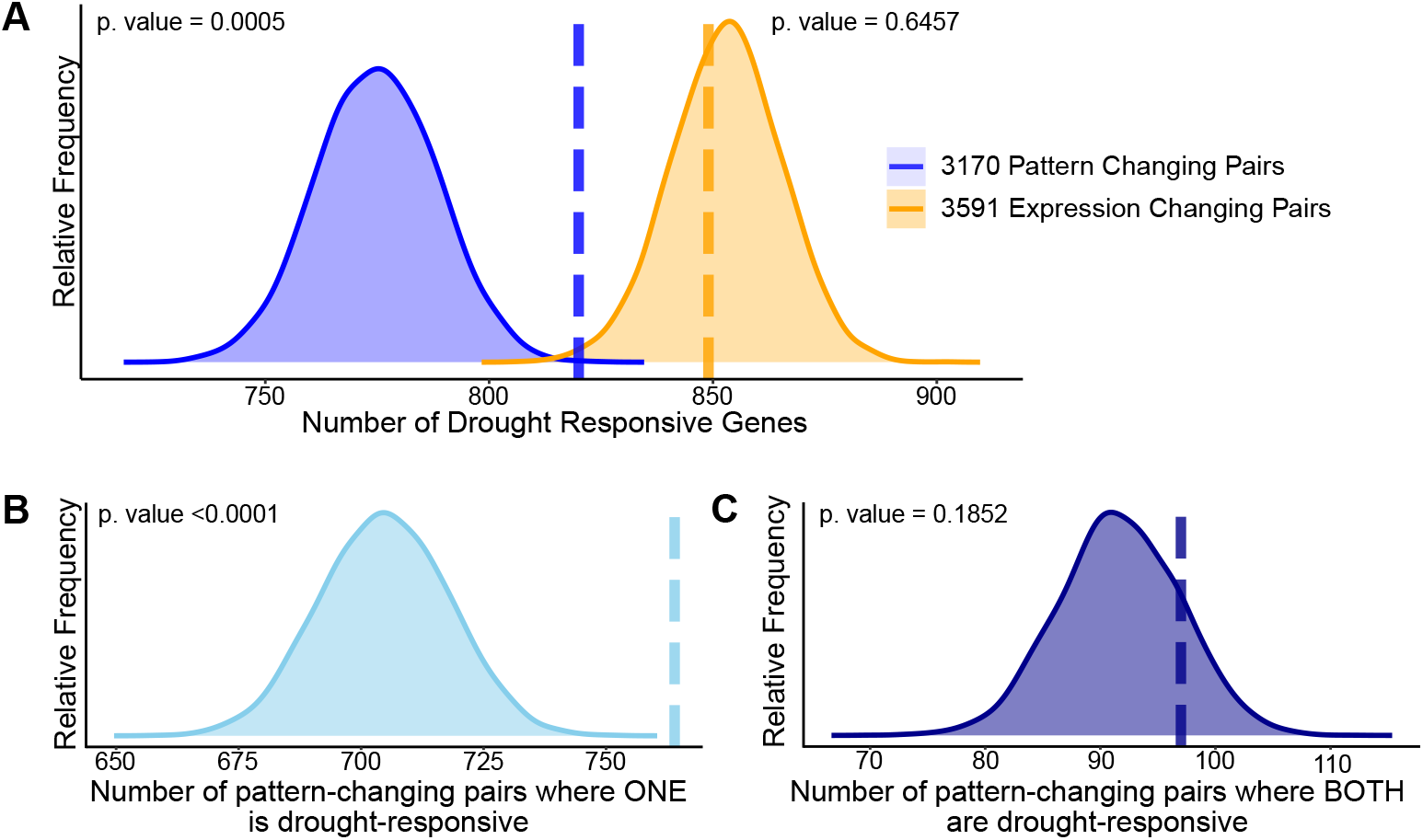
Divergence in drought responsiveness among retained paralogs. (A) Frequency distributions showing the results of a permutation test of the likelihood of paralogous pairs with significantly diverged expression patterns (blue) or significantly different expression levels (orange) in control conditions being drought responsive. Frequency distributions showing permutation results testing the significance of only one member of the paralogous pair being drought-responsive (B) or both paralogs being drought-responsive genes (C). The dashed vertical lines represent the true test statistic and the frequency distributions represent the null distributions that results from the 10,000 permutation tests.

An assessment of functional comparisons among paralog expression levels defines the gene with the highest expression in one or multiple tissue samples as ‘winning’ over the other (3). This classification is often referred to when looking for signs of subgenome dominance within polyploid species. With our set of pairs with one drought responsive paralog, we wondered whether the responsive member of a pair had a higher median expression level under control conditions. Of the 764 pairs with a drought responsive paralog, the drought responsive gene was the lower expressing member in 420 pairs and the higher expressing member in 344 pairs in the control conditions. Thus, we conclude that transcript abundance, whether at a single time point or combined across a time series, is not a reliable predictor of a gene’s functional importance. As validation of this, we ran a standard linear model test at each time point comparing well-watered and drought treatments and did not detect significant transcript abundance differences for the genes identified in the DiPALM analysis. To detect the initial transcriptional response to drought perception we had to incorporate the complete transcript profile into the differential expression test. What appear to be very subtle changes in the abundance of a subset of transcripts are contributing to the measured temporal physiological changes in Fv’/Fm’ and stomatal conductance also occurring at specific time points (26). The ability to capture these early transcriptomic responses to the onset of drought offers new insight into how responsive the network is to slight adjustments in the temporal regulation of expression. The next challenge is to capture this fine-scale resolution across genotypes with diverse physiological responses to stress and identify the associated patterns.

## Discussion

This study emphasizes the power of *B. rapa* as a model system for investigating the consequences of polyploidy on transcriptional network dynamics. The close relationship of *B. rapa* to Arabidopsis facilitates comparative studies and guides gene function hypotheses due to the wealth of genomic and molecular resources developed in Arabidopsis. The development of DiPALM has enabled a new line of inquiry into how temporal regulation of paralogs influence GRNs in *B. rapa*. Replacing single time point comparisons of differential expression with pattern analysis provides a more complete view of the transcriptional network and the pervasiveness of rhythmic gene expression. We have provided further support for the extensive circadian and diel regulation of the transcriptome that has been well documented in Arabidopsis and a few other plant species (22, 43, 44, 46, 47). Our circadian time course experiments with 2h sampling density provided the resolution to reliably assess rhythmicity for all expressed genes resulting in roughly 77% showing circadian clock regulation in *B. rapa* (Fig. 1). The retention of circadian regulation of the transcriptome is consistent with a critical role of the circadian clock in regulating diverse aspects of plant physiology. Interestingly, we found that genes retained in multi-copies in *B. rapa* are enriched in network modules that are phased during the day whereas evening and night phased modules are depleted for multi-copy genes (Fig. 2*D*). This disparity in phase among paralogs might be related to the dosage sensitivity of processes occurring during the day. Gene ontology enrichment processes for the genes in these daytime phased modules included photosynthesis and abiotic stress response (Dataset S3). However, it should be noted that in general, evening phased genes are likely to be greatly understudied because most experiments are performed during the day (45).

The retention of multi-copy genes that are circadian regulated provided an opportunity to assess the level of retention of transcript abundance patterns among paralogs to look for signs of possible neo- or sub-functionalization. Applying DiPALM to our list of circadian regulated paralogs uncovered evidence for extensive rearrangement of the transcriptional network through the divergence in expression pattern among retained paralogs in *B. rapa* (Fig. 3). These changes in phasing among paralogs occur in similar network modules where groups of genes are classified with a particular phase while their paralogous pair-mate exhibit a similar phase difference indicative of common regulatory control. The expansion of expression domains among paralogous pairs provides ample opportunity for neo- and sub-functionalization through new network connections and novel interacting targets now expressed in phase with the pair member with the altered phase of expression. To test this hypothesis and predict the possible network rearrangements that have occurred in *B. rapa* since diverging from Arabidopsis, we took a GRN approach to model the relationships between TFs and gene expression patterns.

Using GENIE3 (34), we input the gene expression pattern information for all TFs and target genes creating a GRN with TFs associated with a group of predicted targets based on expression pattern (Fig. 4*B*). By comparing orthologous TFs between Arabidopsis and *B. rapa* we could assess the overlap in network connections between the TFs and their target genes. This comparison resulted in a set of 49 TF *B. rapa* paralogs where one paralog showed significantly more overlap with the Arabidopsis ortholog (Fig. 4*C*), supporting the divergence in network regulation between retained paralogous TFs in *B. rapa*. This is further supported by the overlap of CNSs among target genes in the GRNs. By replacing genes by CNSs in the GRN we were able to identify the more Arabidopsis-like paralog with strong consensus with the original GRN. The success of the CNS network approach supports a predictive role of these CNSs for expression dynamics and provides a refined nucleotide sequence space to explore in future studies to associate regulatory elements with specific expression patterns. Associating CNSs with specific gene pattern responses may uncover new regulatory elements or novel combinations of regulatory elements that contribute to the differential regulation of paralogs and their targets, for example in response to drought stress. The significant enrichment of one rather than both members of a paralogous pair being drought responsive provides support for possible neo- or sub-functionalization (Fig. 5). That significant amino acid sequence variation apparently does not contribute to the divergence between *B. rapa* paralogous TFs reinforces the importance of regulatory element variation. This raises the question of how two paralogous TFs with the same motif binding affinities can have such diverse targets in the GRN. One possibility is that the presence of new interacting partners at the novel phase of expression could modify binding affinity or prevent binding to certain motifs. Similarly, the lack of critical interacting partners due to a mismatch in phasing could either permit or eliminate some binding targets. Further study into the temporal regulation of known binding partners for the divergent TFs is needed.

These findings bring up several questions surrounding the importance of the variation in paralog expression pattern. Are these differential patterns maintained across *B. rapa* morphotypes or is there additional within-species variation? Did these phase differences arise post genome triplication or were they present in the diploid progenitors that gave rise to *B. rapa*? The morphotypes do exhibit differential circadian clock parameters, as assessed by leaf movement analysis (18), strongly supporting additional within-species variation in the transcriptomic network. An examination of the circadian networks across *B. rapa* morphotypes is needed to characterize these differences and begin to associate network plasticity with morphotype specific traits. Our analysis reveals divergence in drought response among retained paralogs; how do these responses differ in more or less drought tolerant genotypes? Applying these pattern analysis approaches on pan-transcriptome time course studies has the potential to identify regulatory elements that contribute to transcriptional network architecture and the evolution of new forms of transcriptional control in polyploids.

## Materials and Methods

### Circadian Transcriptome Growth Conditions

Seeds of *Brassica rapa* subsp. *trilocularis* (Yellow Sarson) R500 were planted in (3^1/4^” × 3^5/8^”) pots with a soil mixture of 2 parts Metro-Mix PX1 + 1 part Pro-Mix amended with 0.5ml of Osmocote 18-6-12 fertilizer (Scotts, Marysville, OH). The LDHH time course plants were entrained in a 12h light/12h dark cycle at 20°C and the LLHC plants were entrained in a 12h 20°C/12h 10°C under constant light for 15 days. Lights in the chamber at plant height were ~130 μmol photons m^−2^ s^−1^. Plants were shifted to constant light and temperature LLHH for 24h prior to starting the leaf tissue sampling at ZT24. Leaf tissue (~100mg) from the youngest fully developed leaf was harvested and frozen in liquid nitrogen every 2h for 48h (ZT24 – ZT72). At each time point, leaf tissue from 10 plants was collected.

### RNA-sequencing library preparations and processing

Leaf tissue was ground to a fine powder using a Retsch Mixer Mill MM 400 (Vendor Scientific, Newtown PA). The mRNA extraction was performed according to Greenham *et al.* (26) and the strand specific libraries according to Wang *et al.* (48). For each leaf sample (~100mg), 1mL lysis binding buffer (LBB) was used to resuspend ground tissue. For each of two biological replicates, 200μl aliquots of LBB lysate from each of five plants were pooled prior to mRNA isolation. Library size and quality was verified using a 2100-bioanalyzer (Agilent Technologies, Santa Clara, CA). Libraries were indexed and pooled into 12 sample sets and sequenced as 101 bp paired-end reads using Illumina HiSeq2500 (Illumina, San Diego, CA). Raw data have been submitted to GEO (http://ncbi.nlm.nih.gov/geo) under accession number GSE123654. The raw fasta reads were filtered using trimmomatic (49) with mostly default settings (ILLUMINACLIP:./Tru-Seq3-PE.fa:2:30:10 LEADING:3 TRAILING:3 SLIDINGWINDOW:4:25 MINLEN:50). Reads were aligned to the *B. rapa* R500 genome Brapa_R500_V1.2.fasta (https://genomevolution.org/CoGe/OrganismView.pl?org_name=Brassica%20rapa) using tophat2 (https://ccb.jhu.edu/software/tophat/index.shtml) with the following options: --library-type fr-firststrand -I 12000 -G R500_v1.6.gff -M --max-segment-intron 12000 --max-coverage-intron 12000. Sample LD_ZT62_rep2 was identified as an outlier and removed, to avoid over-weighting rep1 the rep2 values were imputed by averaging the values of ZT60_rep1 and ZT64_rep1. Raw counts were generated in Subread version 1.6.3 (http://subread.sourceforge.net/) with the following options: -F SAF -M -T 6 --fraction -s 2 -p -B - Count data was normalized using edgeR (50, 51) (https://bioconductor.org/packages/release/bioc/html/edgeR.html) version 3.22.1 using 'calcNormFactors' and log2 FPKM values were calculated using 'rpkm' with log=TRUE, prior.count=0.1.

### R package DiPALM (differential Pattern Analysis by Linear Model)

We created an R package that takes a raw count table of RNAseq data and runs differential pattern analysis from time series gene expression data. DiPALM is available through the Comprehensive R Archive Network (CRAN) (https://cran.r-project.org/) or via the Greenham Lab Github page (https://github.com/GreenhamLab/Brapa_R500_Circadian_Transcriptome). A sample dataset is provided with the package along with a detailed vignette and manual that describes the analysis pipeline.

### Bioinformatic and Statistical Analysis

The entire analysis pipeline, starting with raw count data, was carried out using the R Statistical Programing Language (27) along with the Rstudio integrated development environment (52). A comprehensive R markdown file is available through the Greenham Lab Github page (https://github.com/GreenhamLab/Brapa_R500_Circadian_Transcriptome). This analysis script includes all data processing, statistical analysis and plotting that was used for this publication. Additional R packages were used in this analysis, including ‘edgeR’ (50, 51), ‘stringr’(53), ‘ggplot2’ (54), ‘rain’ (23), ‘WGCNA’ (25, 55), ‘circlize’ (56) and ‘pheatmap’ (57).

### R500 drought RNAseq dataset

For the drought RNAseq analysis we used our previous dataset (26) and aligned the data to our new R500 genome assembly (https://genomevolution.org/CoGe/OrganismView.pl?org_name=Brassica%20rapa) using the same pipeline described for the circadian datasets. These raw counts are available in a file called ‘DroughtTimeCourse_CountTable.csv’ on the Greenham Lab Github page (https://github.com/GreenhamLab/Brapa_R500_Circadian_Transcriptome).

### CNS Annotation

Using a list of canonical CNS sequences derived from Haudry *et. al.* (39), the R500 and TAIR10 genomes were annotated for those CNSs using NCBI BLAST+. A local BLAST database for each reference genome was created using the command:

~~~
makeblastdb -in <reference.fasta> \
-parse_seqids \
-hash_index \
-blastdb_version 5 \
-dbtype “nucl” \
-title <title>
~~~

To get an initial set of CNS alignments, the following command was run for each reference genome:

~~~
blastn -query <cns.fasta> \
-db <reference.fasta> \
-task “blastn” \
-out <out.csv> \
-outfmt “10 qaccver saccver qlen sstart send sstrand evalue bitscore qcovs” \
-dust “no” \
-soft_masking “false” \
-evalue 0.01 \
-num_threads <num_threads>
~~~

Filtering was disabled in favor of a different scheme also used in Yocca *et. al*.(58). All alignments with a bitscore of 28.2 were dropped, and alignments with a smaller than 60% coverage of the CNS sequence (BLAST+’s “qcovs” value) were also dropped. Finally, to ensure the CNS alignments are reasonably unique, all of the alignments for a particular CNS sequence were discarded if they appeared more than once in the TAIR10 reference, or more than three times in the R500 reference. Since the *B. rapa* genome has undergone a genome triplication event relative to *A. thaliana*, three occurrences were seen as the maximum reasonable amount. Two BED files were generated for each genome containing the coordinates of each resulting alignment.

To associate the resulting CNS alignments with genes in the references, BEDtools was used to find the closest gene to each CNS location:

~~~
bedtools closest -s -t all -D a -a <cns.bed> -b <reference.bed> > CNS_prox_genes.txt
~~~

…where cns.bed is one of the two BED files generated in the previous section, and reference.bed is a gene annotation for the respective reference genome. The -s option constraints reported associations to be only on the same strand -that is, CNS alignments and genes must appear on the same strand. -D a tells BEDtools to report distances, and ensures that the reported distances are signed (negative for occurring before the gene, positive for occurring after, and 0 for being intragenic).

### Motif Analysis

Motif analysis was performed using HOMER --specifically, findMotifsGenome.pl. This program requires that two sets of sequences be provided: a set of target sequences to be searched for motifs, and a set of background sequences for comparison to the target sequences to be compared to. Motif analyses were performed on three different target groups.

The first analysis consisted of searching the CNSs against an “extended” promoter background. For each transcription factor group, the CNSs corresponding to all of the target genes in Ath, Br1, and Br2 separately were pulled and placed into three BED files containing the coordinates of the CNSs in TAIR10 and R500 respectively. A background set of sequences was then generated for every gene in the TAIR10 and R500 genomes using BEDtools:

~~~
bedtools slop -s -i <reference.bed> -g <reference.genome> -l 2000 -r 0 > 2kb_and_gene.bed
~~~

…where reference.bed is a gene annotation for the reference genome, and reference.genome is a text file containing the lengths of each chromosome that BEDtools uses to ensure that the coordinates it outputs are valid. This outputs a BED file that annotates a background consisting of a 2kb promoter region before each gene, as well as the gene itself, to the end of the 3’ UTR. This larger background sequence was selected rather than the 2kb promoter alone since many CNSs occurred in the UTRs and introns, and so using only 2kb promoters would leave out background sequence relevant to many of the CNSs, potentially skewing the results.

The second analysis searched the “extended” promoter sequences against themselves. The target set consisted of “extended” promoters corresponding to target genes in each transcription factor group, and the background consisted of a total list of sequences, including the target group, as per HOMER’s recommendations.

The third analysis consisted of a more traditional motif search of promoter regions against promoter-only background. In this case, neither the target nor background sequences contain CDS, introns, or UTRs as with the “extended” regions defined in the previous two analyses. These promoters were pulled using BEDtools:

~~~
bedtools flank -s -i <reference.bed> -g <reference.genome> -l 2000 -r 0 > 2kb_promoters.bed
~~~

Everything is identical as above, except that bedtools flank does not include the genes themselves in the output, and only generates locations for promoter regions. As before, promoters corresponding to TF target groups were selected and then analyzed against the entire set of promoters. HOMER was run using its included plant motif database on all three datasets (Datasets S7-S8). Default parameters were used.

Motif analyses were performed separately for the Ath, Br1, and Br2 target groups. Given that three analyses were performed for each of these target groups, a total of 9 motif analyses were performed for each TF group for a total of 108 motif analyses. Only the “knownMotifs” output of HOMER was considered, which consists of a database search of target and background sequences against known plant motifs with a hypergeometric test to quantify significance. The de novo results were not used. For each of the 108 analyses, the outputs of the “knownMotifs” analysis were simplified by grouping together found motifs that correspond to the same DNA-binding protein domain. Of each of these domain groups, the best p-value out of all the motifs found for that domain were selected as representative for the entire group. For each TF group, significant (p<0.01) domains were counted for the Ath, Br1, and Br2 target groups, for the CNS, “extended” promoter, and promoter-only motif analyses.

## Author contributions

K.G., T.C.M., and C.R.M. designed research; K.G. and P.L. performed experiments; K.G., R.C.S., and S.Z. analyzed data; and K.G., R.C.S., and C.R.M. wrote the paper.

## Conflict of Interest

The authors declare no conflict of interest.

## Acknowledgements

This work was supported by National Science Foundation grants IOS-1202779 to K.G., IOS-1711662 to R.S., IOS-1547796 and by the Rural Development Administration, Republic of Korea Next Generation BioGreen. 21, grant number SSAC PJ01327306 to C.R.M.

