## Supplemental Figures S1-S3 for "Expansion of the circadian transcriptome in *Brassica rapa* and genome-wide diversification of paralog expression patterns"

**Supplementary Material**
Figures S1 to S3

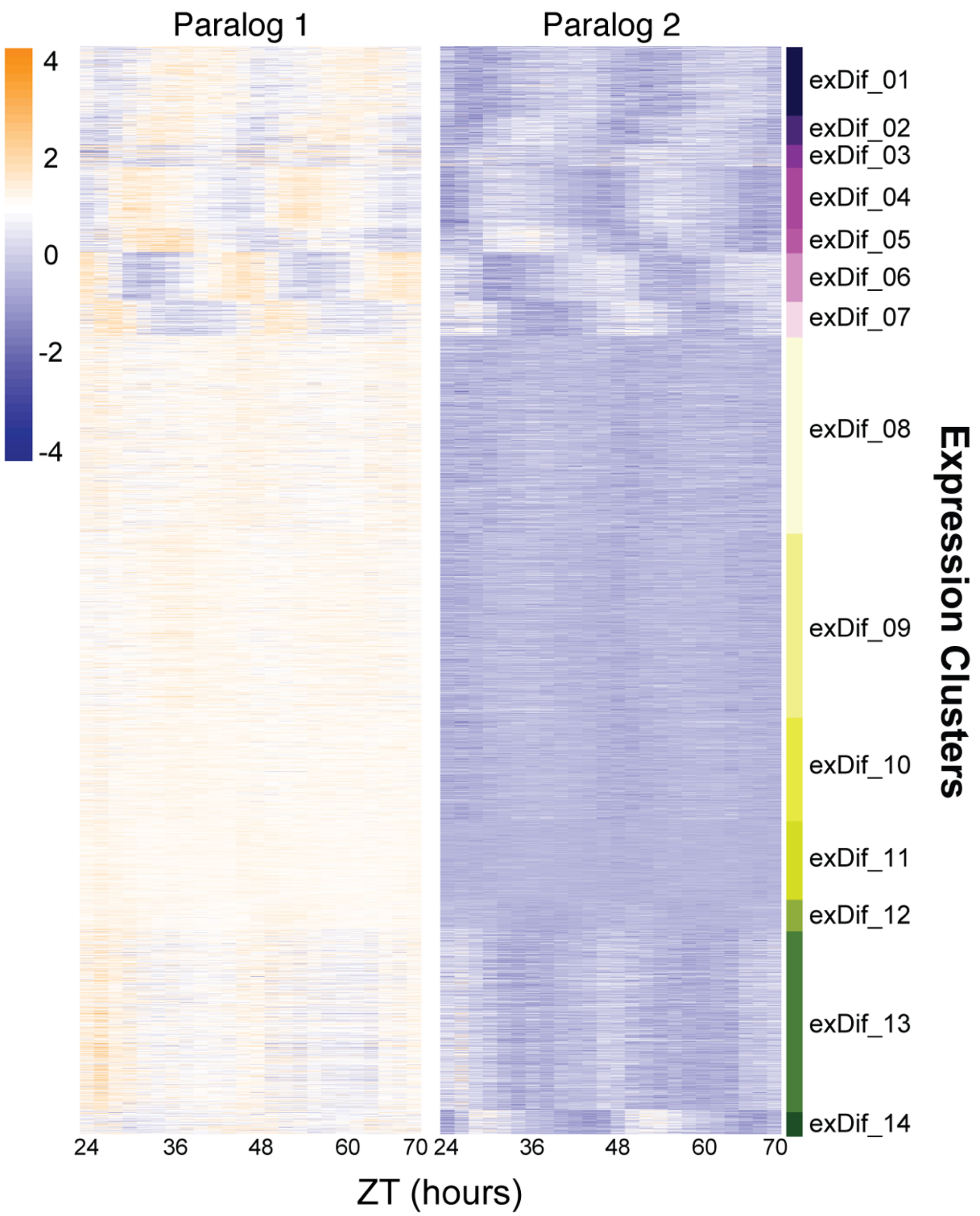


**Figure S1 Divergence in median expression levels among retained paralogs.** Heat map of the results from the DiPALM exDif clustering showing the changes in expression pattern for paralogous pairs. Each line of the heat map for each block corresponds to a paralogous pair. Three copy paralogs were split into three, 2-way comparisons. Expression values are log2 transformed FPKM values and the expression is arranged by ZT time across the x-axis. Higher expression levels are orange and low expression levels are purple.

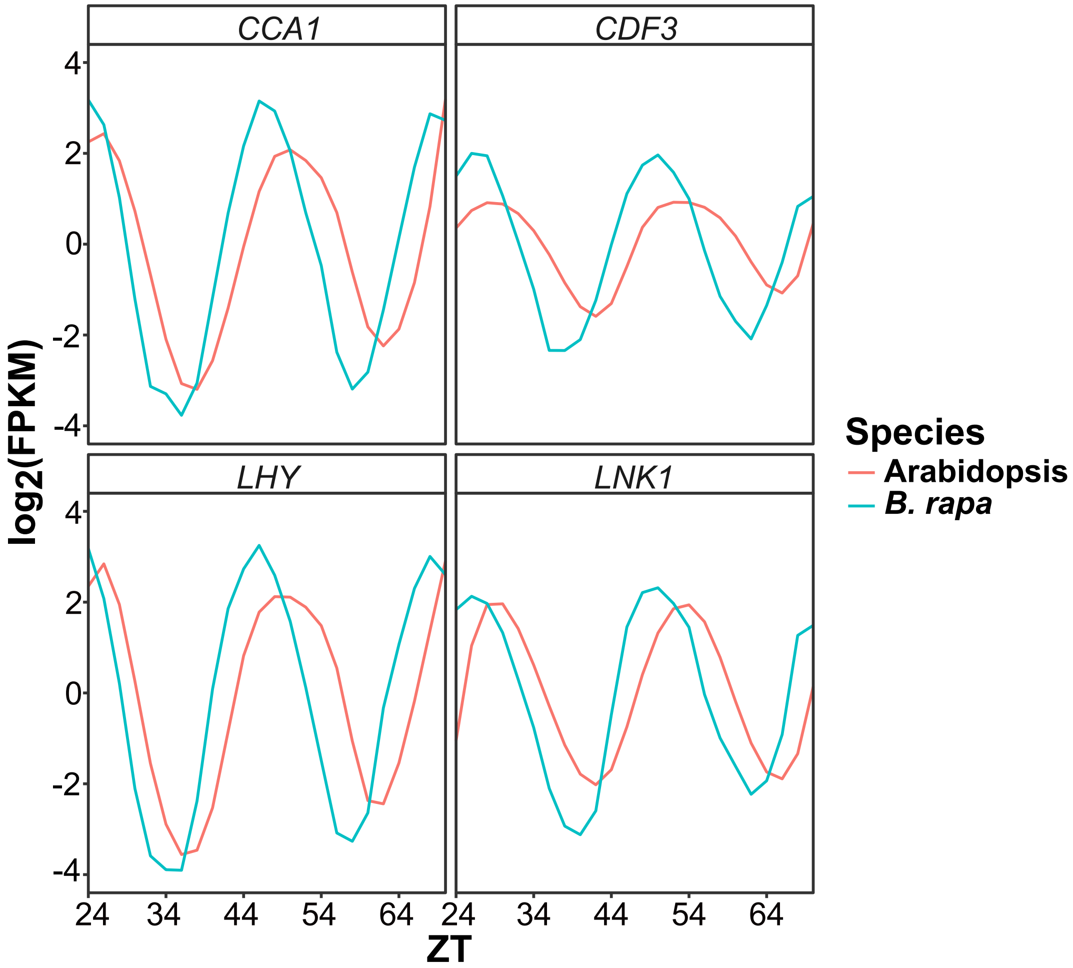


**Figure S2 Phase variation between Arabidopsis and *B. rapa* clock genes.** Gene expression expressed as log2 transformed FPKM values for four circadian clock genes (*CCA1, CDF3, LHY*, and *LNK1)* from the Arabidopsis microarray data and *B. rapa* RNAseq data used in this study. Due to the variation in circadian period in Col-0 and R500 there are visible phase differences as seen by the shift in peak timing of expression.

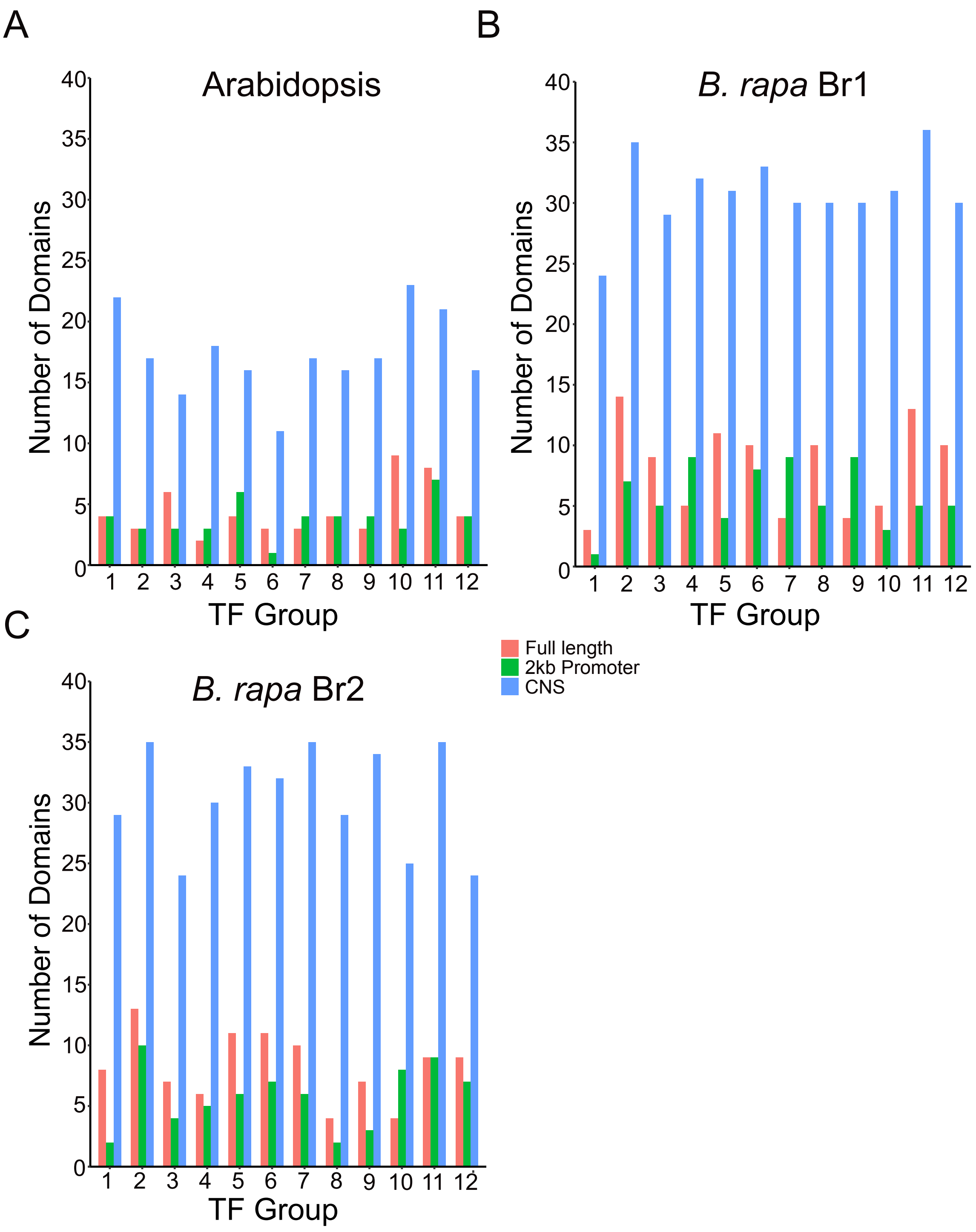


**Figure S3 Motif enrichment in CNS regions in Arabidopsis and *B. rapa* GRN TF target genes.** The HOMER motif analysis software was used to look for enrichment of known TF binding motif sequences among the target gene groups for the top 12 TFs in the gene and CNS GRN overlap list (Figure 4G). Motif results were compared for the CNS regions, the full-length gene (2kb upstream promoter to the 3' UTR) and just the 2kb promoter region (2kb upstream promoter to the start codon). For the Arabidopsis (A), Br1 (B) and Br2 (C) target group analysis there was between 3-5 fold greater enrichment of motifs in the CNS regions.
